# Cerebrospinal fluid outflow through skull channels instructs cranial hematopoiesis

**DOI:** 10.1101/2021.08.27.457954

**Authors:** Fadi E. Pulous, Jean C. Cruz-Hernández, Chongbo Yang, Zeynep Kaya, Gregory Wojtkiewicz, Diane Capen, Dennis Brown, Juwell W. Wu, Claudio Vinegoni, Masahiro Yamazoe, Jana Grune, Maximillian J. Schloss, David Rohde, Dmitry Richter, Cameron S. McAlpine, Peter Panizzi, Ralph Weissleder, Dong-Eog Kim, Filip K. Swirski, Charles P. Lin, Michael A. Moskowitz, Matthias Nahrendorf

## Abstract

Interactions between the immune and central nervous systems strongly influence brain health. Although the blood-brain barrier restricts this crosstalk, we now know that meningeal gateways through brain border tissues, particularly dural lymphatic vessels that allow cerebrospinal fluid outflow, facilitate intersystem communication. Here we observe that cerebrospinal fluid exits into the skull bone marrow. Fluorescent tracers injected into the cisterna magna of mice travel through hundreds of sub-millimeter skull channels into the calvarial marrow. During meningitis, bacteria usurp this perivascular route to infect the skull’s hematopoietic niches and initiate cranial hematopoiesis ahead of remote tibial sites. Because skull channels also directly provide leukocytes to meninges, the privileged sampling of brain-derived danger signals in cerebrospinal fluid by regional marrow has broad implications for neurological disorders.

**One-Sentence Summary:** Skull channels transport cerebrospinal fluid from the subarachnoid space to the cranial bone marrow via a perivascular route, which bacteria use during meningitis.

## Introduction

In addition to guarding brain health, the immune system participates in a wide array of neurological disorders. The blood-brain barrier enforces an unusually rigid leukocyte origin dichotomy, dividing central nervous system (CNS)-resident immune cells from systemically circulating leukocytes. Crosstalk between local brain and systemic immune system components is limited in the steady state but expands during pathologies. Residing at potential portals of entry, the meninges are CNS border tissues that provide the brain and spinal cord with a protective connective tissue capsule consisting of three tissue layers (pia mater, arachnoid and dura mater) within which cerebrospinal fluid (CSF) flows between the arachnoidal membrane and pia mater. A complementary glymphatic system (*1*), which drains the brain’s interstitial and perivascular spaces, interfaces with CSF. Ultimately, CSF exits into the venous blood and the lymphatics (*2, 3*).

The CNS border tissues dynamically police leukocyte migration and brain-derived signal exit. Recently discovered skull channels connecting the cranial bone marrow to the meninges, in mice and humans, constitute a novel leukocyte portal into the CNS (*4–6*). Skull hematopoietic activity directly adjacent to the brain delivers myeloid cells (*4, 6*) and B lymphocytes (*5*), bypassing the blood-brain barrier. If the skull marrow is indeed a private leukocyte purveyor for the brain, skull-derived immune cells may be an intermediate third entity between resident and non-resident CNS leukocytes. Further, dural lymphatics provide a CSF outflow which freely exchanges with the brain’s glymphatic system (*1*). This signaling pathway shares information about brain health systemically, allowing for presentation of CNS-derived antigen in cervical lymph nodes and consequently activating adaptive immunity (*7, 8*). Collectively, these reports expand our understanding of immune cell function across CNS borders and specifically implicate CSF as an under-appreciated messenger that may coordinate neuroinflammation.

Here we describe previously unrecognized CSF outflow into the skull bone marrow. Fluorescent tracers injected into the cisterna magna of mice migrate along the perivascular spaces of dural blood vessels and then perivascularly travel through skull channels into the cranial marrow. In mice with meningitis, bacteria usurp this path into the skull marrow, thereby boosting cranial emergency hematopoiesis.

## Results

### Perivascular CSF transit through skull channels into marrow cavities

To understand the spatial organization of the skull channels, we first performed high-resolution *ex vivo* X-ray computed tomography of the skull (Fig. 1A) and characterized regional channel networks overlying frontal, parietal and occipital brain lobes (Fig. 1B). Channels traversed the inner compact bone into the marrow-containing cavities. We observed the highest density of skull channels in the frontal and occipital regions (Fig. 1C). Given a CT-derived mean channel density above 10 per mm^2^ and an inner skull surface area >100 mm^2^, we estimate that more than 1,000 channels reach into the cranial vault of an adult mouse. Frontal and parietal skull channels formed the shortest connections to the dura, ranging from 83–90 μm, whereas occipital skull channels were approximately 25% longer (Fig. 1D). Frontal and parietal skull channels were 20% narrower than their occipital counterparts (Fig. 1E), pointing to regional channel heterogeneity.

**Fig. 1.**
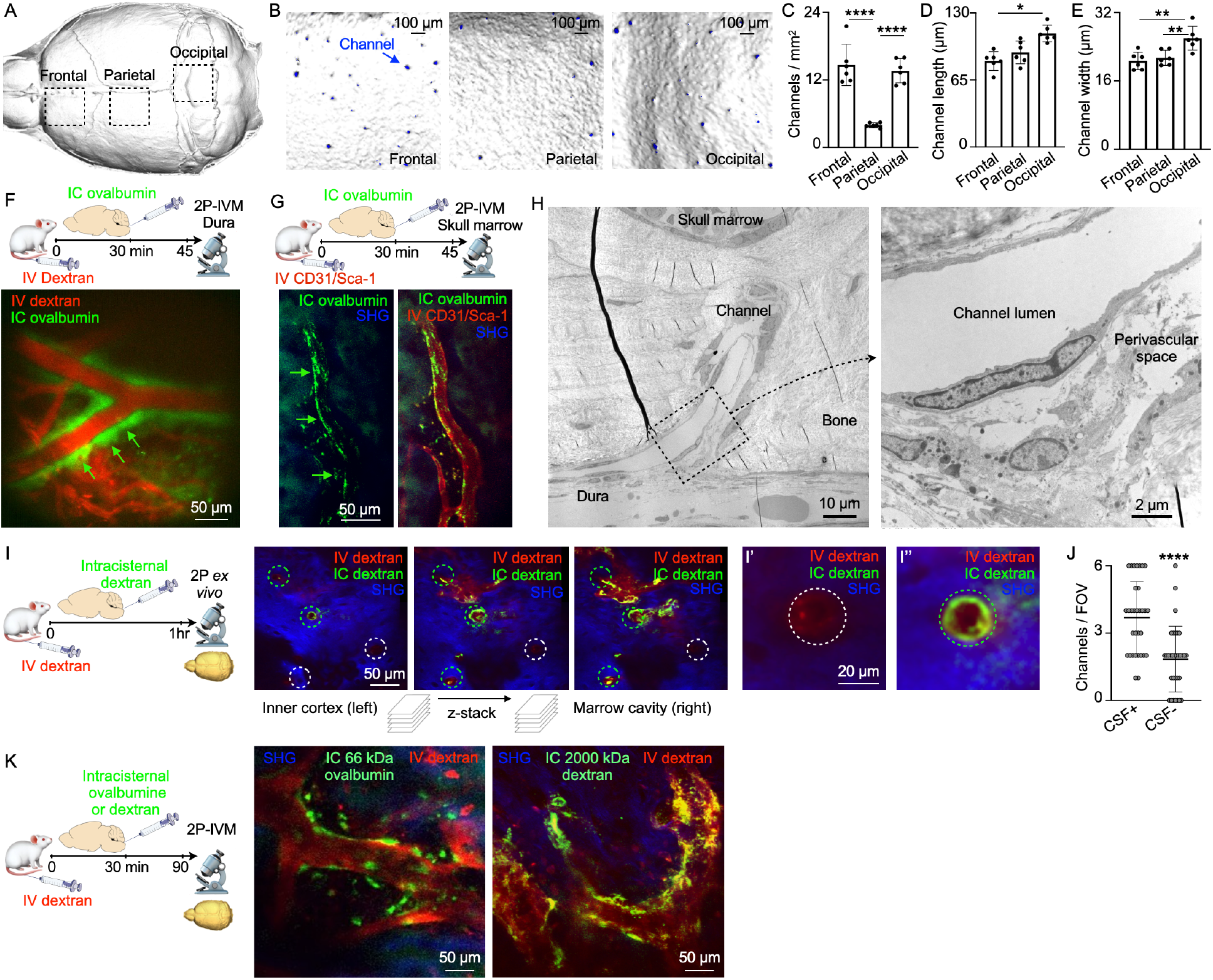
A perivascular space in skull channels facilitates cerebrospinal fluid outflow into the cranial marrow. (**A**) Inner skull cortex 3D surface reconstruction from microCT. (**B**) 3D reconstructions of inner frontal, parietal and occipital bone surfaces with channel openings indicated in blue(scale bar: 100 μm). Regional channel density (**C**), length (**D**) and width (**E**) measured by microCT (mean ± SD; n=6 mice; one-way ANOVA with Tukey’s multiple comparisons test; \**P*<.05, \*\**P*<.01, \*\*\*\**P*<.0001). (**F**) Scheme and intravital microscopy (IVM) image of intracisternally (IC) injected ovalbumin in the perivascular space of a dural vessel intravenously (IV) labeled with dextran (scale: 50 μm). (**G**) Scheme and IVM image of intracisternally injected ovalbumin in the perivascular space of a bone marrow vessel (scale: 50 μm). (**H**) Transmission electron microscopy of a skull channel in the frontal bone (scale: 10 μm). Inset of perivascular area (scale: 2 μm). (**I**) Z-stack of interior skull cortex *ex vivo* microscopy in mice after intracisternal and intravenous injections of dextran. Bone signal from second harmonic generation surrounds channels (circles). I’ and I’’ are insets of IC tracer negative and positive channels (scale: 50 μm and 20 μm). (**J**) Number of CSF-containing channels (mean ± SD; n=3 mice; Mann-Whitney test; *P*<.0001). (**K**) IVM of mice after intracisternal injection of ovalbumin/dextran for CSF tracing and IV dextran (scale: 50 μm, n=3-5 mice per tracer; scale: 50 μm).

We hypothesized that these numerous links between the skull marrow and the dura enable not only cell traffic towards the meninges (*4*) but also bidirectional crosstalk. We therefore implemented a 2-photon intravital microscopy (IVM) and *ex vivo* imaging pipeline to visualize CSF distribution after fluorescent tracer injection into the cisterna magna (*9, 10*). Intracisternal injection of 70 kD FITC-labeled ovalbumin was combined with intravenous labeling of the blood pool using 70 kD Texas red-labeled dextran. We observed a striking perivascular appearance of the intracisternal tracer (Movie S1 and Fig. 1F) along a subset of dural vessels, indicating that CSF travels perivascularly from the subarachnoid space into the dura. This led us to wonder whether we could track this CSF outflow into the skull marrow cavity. We labeled cranial vasculature with intravenously injected fluorescently conjugated CD31/Sca1 antibodies and CSF with intracisternally injected fluorescently labeled ovalbumin, followed by intravital imaging of the skull marrow cavity (*4, 11, 12*). In the skull marrow, we detected perivascular spaces and cells labeled with the intracisternally injected CSF tracer (Fig. 1G). This critical observation in the dura and marrow suggested that perivascular CSF outflow into the marrow may occur through skull channels.

As we anticipated, high magnification transmission electron microscopy analysis of skull channels revealed a perivascular space that may accommodate CSF transport beginning at the dural channel opening (Fig. 1H). To directly test this hypothesis, we labeled cranial vasculature with intravenous Texas red dextran and the CSF with an intracisternal FITC dextran injection one hour prior to *ex vivo* microscopy of the inner skull cortex. Channel cross sections were clearly demarcated by dextran-labeled vessels surrounded by bone visualized with second harmonic generation (Fig. 1I). Z-stacks that began on the dural surface and moved deep into the marrow cavity (Fig. 1I and Movie S2) visualized intracisternally injected dextran in numerous skull channels. This signal surrounded the blood vessel and was present from channels’ dural openings all the way into the marrow cavity. Counting CSF-tracer-containing channels revealed that 67% of them showed perivascular signal after intracisternal dextran injection (Fig. 1J). Bone marrow imaging after intracisternal injection revealed a similar perivascular appearance for tracers with molecular weights from 66kD to 2000 kD along a subset of skull marrow vessels (Fig. 1K). Labeled CSF was detected in the skull marrow as early as 15-30 minutes after injection but was largely absent from the tibia’s marrow vasculature (Fig. 1G and S1); this indicates that CSF was excluded from systemic circulation at early time points. Together, these data demonstrate that CSF exits the subarachnoid space via perivascular flux along a subset of dural vessels which connect into the bone marrow cavity through an extensive skull channel network. This finding implicates the skull marrow as a CSF-sensing hematopoietic compartment. We next sought to examine these observations’ relevance in a mouse model of bacterial meningitis.

### Streptococcus pneumoniae *expand near dural skull channel openings*

We adapted a model of pneumococcal meningitis (*13*) to test the functional significance of skull channel connections in neuroinflammation. *Streptococcus pneumoniae* is the clinically dominant cause of bacterial meningitis (*14*). To establish a disease timeline in mice, we injected 5 × 10^3^ bioluminescent *Streptococcus pneumoniae* Xen10 bacteria into the cisterna magna and analyzed bacterial propagation over time alongside control mice that received an equal volume of artificial CSF (Fig. 2A). Whole-body bioluminescence (BLI) imaging revealed a time-dependent signal increase reporting bacterial growth (Fig. 2B). BLI signal was observed predominantly in the skull 36 hours after injection and by 48 hours had spread to the spine. Bacterial burden increased exponentially by 36 hours, with a peak at 48 hours after injection (Fig. 2C). To assess meningeal inflammation, we measured the canonical inflammatory cytokines *Il1β*, *Il6* and *TNFα* in the meninges by qPCR and found them to be 40- to 60-fold higher in mice with meningitis relative to controls injected with artificial CSF (Fig. S2). We next assessed *S. pneumoniae* growth in the blood and CSF using a bacteria colony forming unit assay 48 hours after infection. The blood contained a miniscule amount of *S. pneumoniae*, while the CSF contained approximately 10,000-fold more. This suggests that 48 hours after infection, bacterial propagation is mostly confined to the meninges (Fig. 2D). We next sought to visualize *S. pneumoniae* manifestation relative to the skull marrow cavity and skull-dural channel connections. To this end, we injected green fluorescent protein (GFP)-expressing *S. pneumoniae* strain D39V hlpA-GFP into the cisterna magna and adapted an optical clearing protocol (*15*) to show skull channels, the marrow cavity and adjacent bacterial propagation in the subarachnoidal space. The skull marrow vasculature was stained with intravenously injected fluorescent CD31/Sca-1 antibodies and the bone with osteosense, allowing us to identify intact marrow, channels and the CSF space by confocal microscopy (Fig. 2E). To visualize GFP^+^ *S. pneumoniae*, we employed the timeline established by bioluminescence analysis (Fig. 2A-C). We injected 5 × 10^3^ GFP^+^ *S. pneumoniae*, or an equal volume of artificial CSF in controls, and sacrificed mice 48 hours later (Fig. 2F). Confocal microscopy of cleared tissue revealed abundant GFP^+^ bacterial growth in the subarachnoid space of mice with *S. pneumoniae* meningitis, but not controls (Fig. 2E, G and Movie S3). Three-dimensional reconstructions document the proximity of GFP^+^ *S. pneumoniae* to skull channels (Fig. 2H and Movie S4). In addition to subarachnoidal bacterial colonies directly adjacent to dural skull channel openings, a smaller number of bacteria were also present in the skull marrow’s extravascular space (Fig. 2H). Taken together, these findings gave rise to the hypothesis that in meningitis, bacteria may enter the skull bone marrow. While we know skull fractures may cause bacterial meningitis, intact skull invasion from within the cranial vault has not been explored previously.

**Fig. 2.**
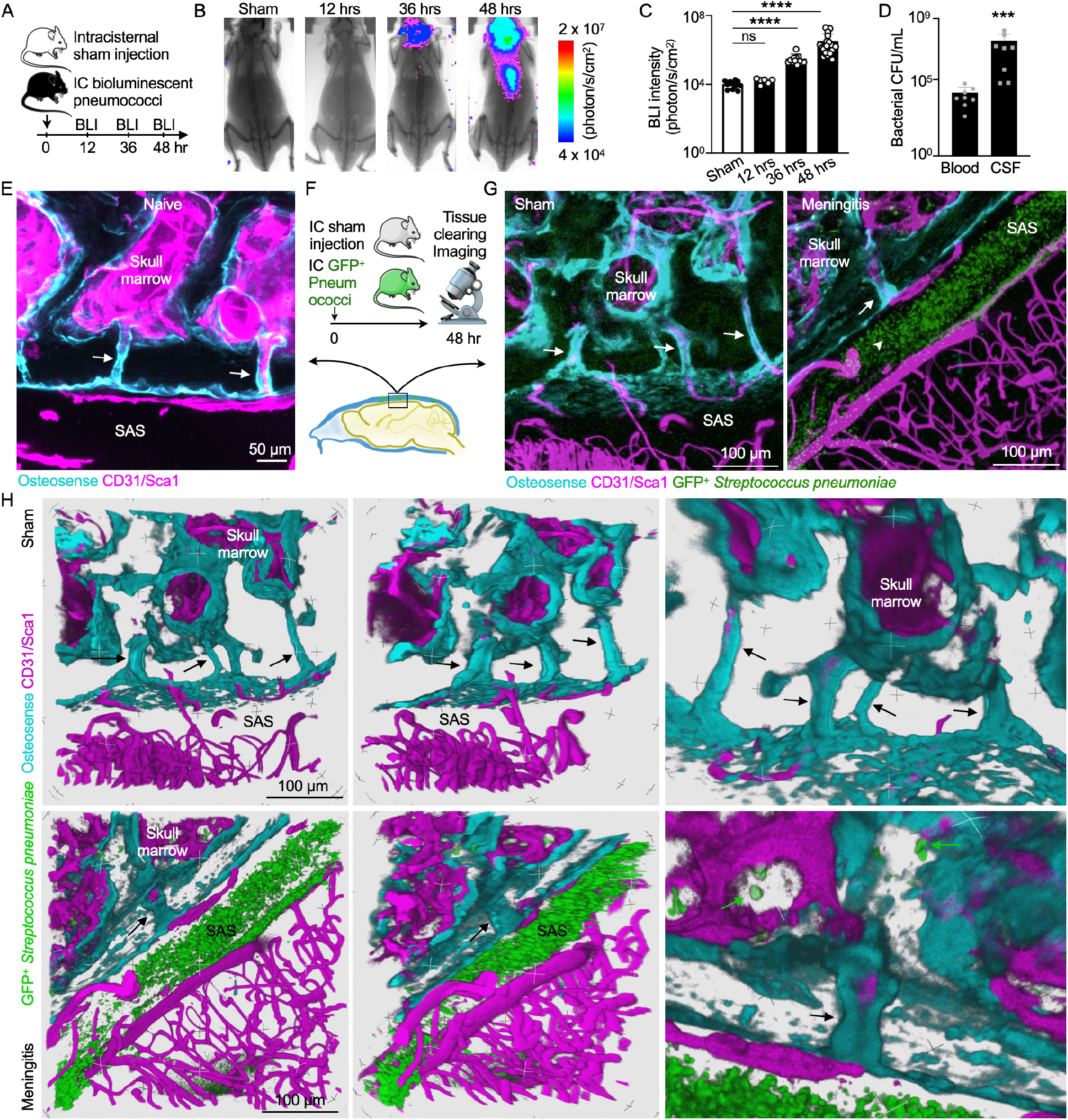
Bacterial propagation during pneumococcal meningitis. (**A**) Timeline for bioluminescent *Streptococcus pneumoniae* Xen10 meningitis. (**B**) Bioluminescence imaging (BLI) of sham controls or mice after intracisternal injection of *S. pneumoniae* Xen10 at indicated timepoints. (**C**) Quantitation of bacterial load with BLI (mean ± SD; n=6-24 mice; Kruskal-Wallis test with Dunn’s multiple comparisons test; ns=not significant, \**P*<.05, \*\*\*\**P*<.0001). (**D**) Bacterial colony forming unit (CFU) assay from blood and CSF 48 hrs after infection (mean ± SD; n=8 mice; Mann-Whitney Test; \*\*\**P*<.001). (**E**) Skull channels visualized after tissue-clearing (scale: 50 μm). (**F**) Experimental outline for tissue clearing and meningitis imaging. (**G**) Representative images of sham controls or mice after intracisternal injection of *Streptococcus pneumoniae* JWV500. Osteosense identified bone while IV CD31-Sca1 labeled BM and dural vessels (n=4-6 mice; scale: 100 μm). (**H**) 3D reconstructions of (G) highlighting skull channels (arrows) and bacterial propagation in marrow (scale: 100 μm).

### Pneumococcal meningitis propagates to the skull

To test the hypothesis that *S. pneumoniae* enter the skull marrow cavity, we performed *in vivo* confocal microscopy of the intact skull in mice with meningitis. Mice were imaged 48 hours after intracisternal injection with GFP^+^ *S. pneumoniae* or artificial CSF in controls. The vasculature was labeled with intravenous CD31/Sca-1 antibodies one hour prior to imaging and the bone was labeled via osteosense injection 24 hours prior to imaging. Surprisingly, we observed GFP^+^ bacterial colonies in the skull marrow extravascular spaces of mice with meningitis (Fig. 3A, Movie S6). No such signal was detectable in controls (Fig. 3A and Movie S5). Since this, to our knowledge, is the first observation of *S. pneumoniae* entering the skull cavity during meningitis, we sought to corroborate these imaging data with orthogonal assays, including bacterial cultures, qPCR for bacterial genes and flow cytometry to detect GFP expressed by bacteria.

**Fig. 3.**
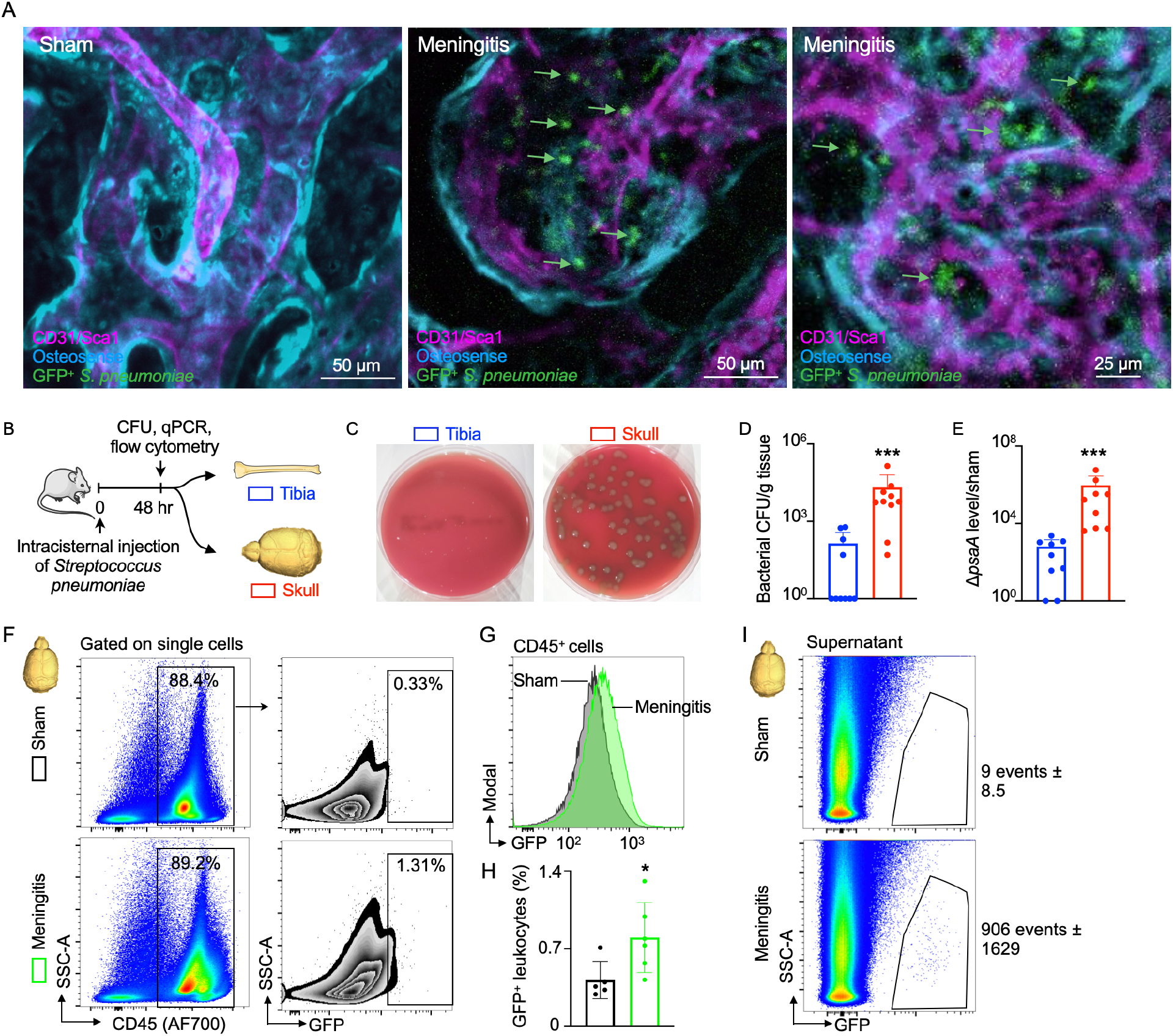
Bacterial migration to skull marrow. (**A**) IVM of skull marrow from sham controls injected with artificial CSF (aCSF) or mice 48 hrs after intracisternal injection of GFP^+^ *Streptococcus pneumoniae* JWV500 show bacterial presence adjacent to vasculature within the bone marrow (n=3-4 mice per group; scale: 50 μm, scale: 25 μm). (**B**) Experimental outline for experiments in (C-G). (**C**) Bacterial cultures from mice with pneumococcal meningitis relative to sham controls. (**D**) Quantitation of bacterial colony forming unit (CFU) assay in (C) shows minimal bacteria in the tibia relative to the skull (mean ± SD; n=10 per group; Mann-Whitney test; \*\*\**P*<.001). (**E**) qPCR of *S. pneumoniae* surface adhesion gene (*psaA*) expression in tibia versus skull marrow normalized to sham controls (mean ± SD; n=9 mice; Mann-Whitney test; \*\*\*\**P*<.0001). (**F**) Gating strategy for CD45^+^ leukocytes that have interacted with GFP^+^ *S. pneumoniae*. (**G**) Histogram of GFP signal in CD45^+^ leukocytes obtained from mice with meningitis compared to CD45^+^ cells from sham-injected controls. (**H**) Quantitation of GFP^+^CD45^+^ cells analyzed in (F,G) (mean ± SD; n=5 mice; unpaired t-test; \**P*<.05) (**I**) Extracellular GFP^+^ *S. pneumoniae* detected by flow cytometry in supernatant of skull marrow from mice with meningitis compared to sham controls (n=2-4 mice per group).

Using an experimental timeline (Fig. 3B) comparable to the microscopy experiments described above, we first employed a bacterial colony forming unit (CFU) assay to analyze bacterial growth 48 hours after intracisternal injection of *S. pneumoniae*. Tibia and skull bone marrow was harvested, homogenized and plated on blood agar plates to accommodate bacterial colony growth, similar to clinical blood cultures. While bacteria were nearly undetectable in tibial marrow, bacterial colonies grew from the skull marrow preparations (Fig. 3C, D). Since only viable bacteria can divide, this finding documents that live bacteria were present in the skulls of mice with meningitis. We next compared skull and tibial marrow using qPCR analysis for the bacterial gene *psaA*, which is not expressed in mice. Skull samples from mice with meningitis contained markedly higher levels of *psaA* transcript compared to the tibia (Fig. 3E), confirming the presence of *S. pneumoniae* within the skull marrow. Furthermore, we performed flow cytometric analysis on skull marrow isolated from mice after intracisternal injection of GFP^+^ *S. pneumoniae*. In addition to detecting bacterial presence, flow cytometry also determined whether bacteria are located inside cells. The skulls of mice with meningitis showed substantial numbers of CD45^+^ GFP^+^ leukocytes, which were largely absent in the skulls of control mice (Fig. 3F). We document the observed fraction of bacteria-containing leukocytes in a right-shifted GFP histogram in CD45^+^ leukocytes obtained from the skull marrow of mice with meningitis, as compared to controls injected with artificial CSF (Fig. 3G and 3H). We also analyzed the cell-free supernatant of bone marrow suspensions following high-speed centrifugation by flow cytometry. In the skull marrow supernatant obtained from mice that received intracisternal GFP^+^ *S. pneumoniae* injection, we noted abundant bacteria that were absent in controls (Fig. 3I). Taken together, the imaging observation of bacteria in the skull marrow of mice with meningitis was confirmed by three independent assays, all supporting that bacteria can propagate from the meninges to the skull marrow.

### Skull channels are conduits for *S. pneumoniae* to access the calvarial marrow

We next sought to directly evaluate whether *S. pneumoniae* reach the skull marrow by transiting skull channels from the dura. Accordingly, we performed *ex vivo* confocal microscopy of tissue-cleared skull preparations containing intact brain tissue 48 hours after intracisternal injection of either GFP^+^ *S. pneumoniae* or artificial CSF in controls. Z-stack projections of skull channels revealed the presence of GFP^+^ *S. pneumoniae* both in the dura and inside skull channels (Fig. 4A). We found that 75% of mice with meningitis showed bacterial GFP signal in their skull channels, whereas control animals without meningitis lacked any sign of *S. pneumoniae* (Fig. 4B). Three-dimensional reconstructions of skull channels revealed extravascular GFP localization within the channels of mice with meningitis, but not in controls (Fig. 4C and Movies S7 and S8). As a complementary approach to imaging bacterial GFP directly, we adapted the CUBIC method to search for GFP^+^ *S. pneumoniae* in deeper tissue areas (*15*). More specifically, we modified the CUBIC protocol II (*15*) to include delipidation, decolorization, decalcification, refractive-index matching and ultimately immunostaining for bacterial GFP with a fluorescently labeled anti-GFP antibody. Whole-mount confocal microscopy of CUBIC-processed specimens revealed a striking pattern of anti-GFP staining within skull channels and in the dura mater (Fig. 4D). Skulls from control mice showed no GFP signal in the dura, channels or marrow (Fig. 4D). Collectively, these data demonstrate two critical and previously unknown phenomena: i) during meningitis, bacteria enter the skull marrow through skull channels from the dura, and ii) bacterial influx through channels into the marrow likely occurs via a perivascular route, similar to CSF outflow into the marrow as observed in the steady state.

**Fig. 4.**
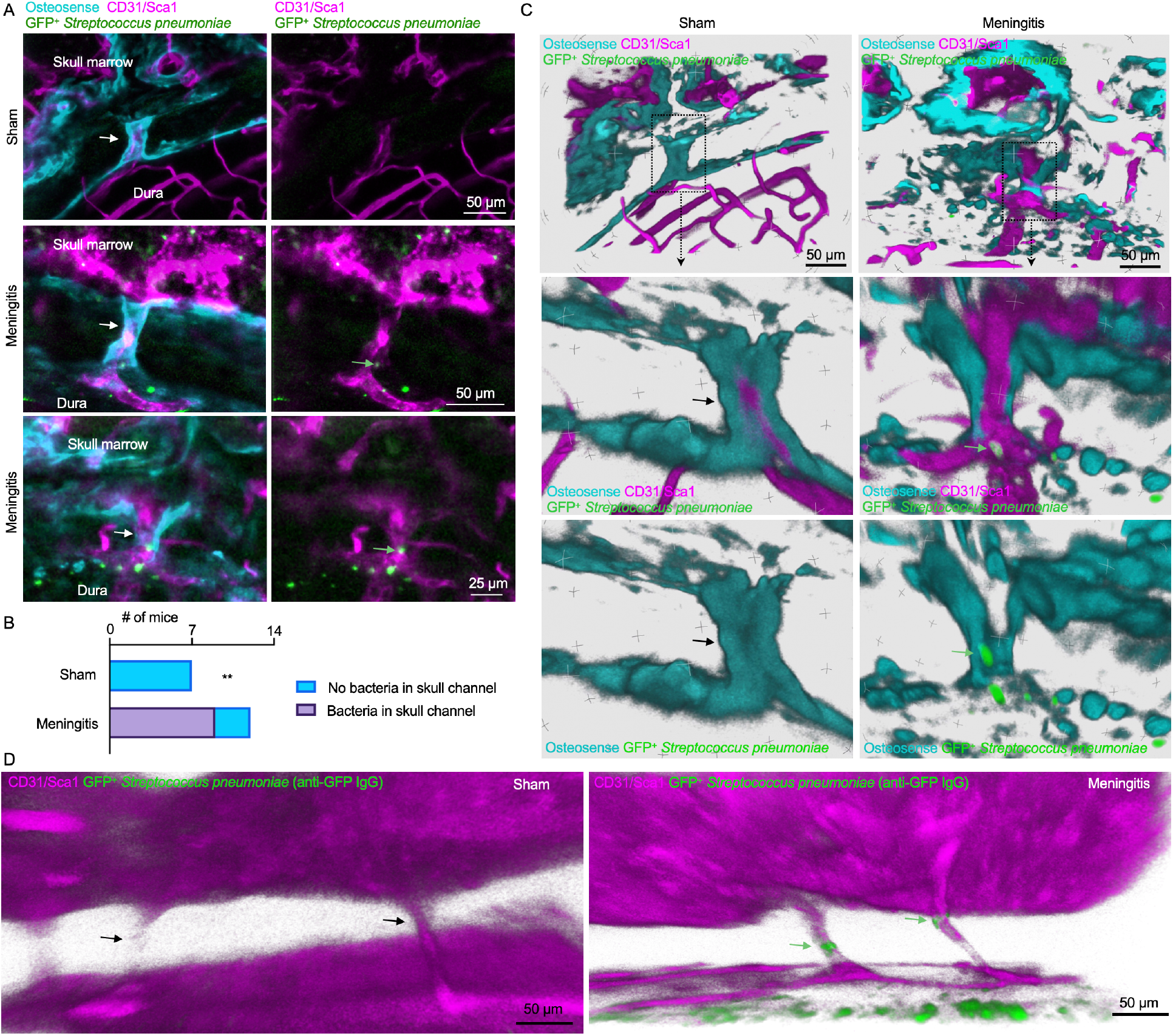
Skull channels are conduits for pneumococcal migration into the cranial marrow. (**A**) Whole-mount *ex vivo* imaging of skull channels in sham controls or mice after intracisternal injection of GFP^+^ *Streptococcus pneumoniae* JWV500 after tissue clearing. Images depict bacteria in skull channels which are visualized using osteosense to label bone and marrow vasculature using a CD31-Sca1 cocktail (n=7-11 mice per group; scale: 50 μm or 25 μm as indicated). (**B**) Quantitation of *S. pneumoniae* GFP signal in skull channels in sham controls and mice with meningitis (n=7 controls, n=11 mice with meningitis; Chi-Square Test; \*\**P*<.01). (**C**) 3D renderings of (A) show intra-channel perivascular localization of *S. pneumoniae* (scale: 50 μm). (**D**) Whole-mount *ex vivo* imaging after CUBIC tissue processing for GFP detection after intracisternal injection of GFP^+^ *Streptococcus pneumoniae* JWV500. CUBIC protocol (described in methods) was followed by immunostaining for bacterial GFP. Bone marrow vasculature was labeled *in vivo* with CD31-Sca1 (n=2 mice; scale: 50 μm).

### *S. pneumoniae* propagation within the CSF induces a skull-specific increase in LSK proliferation

Having defined the route by which *S. pneumoniae* transit skull channels via perivascular passage and arrive within the skull marrow, we next tested how this bacterial expansion functionally affects hematopoiesis. Specifically, we devised an experimental timeline to test whether intracisternal injection of *S. pneumoniae* elicits a hematopoietic response in skull marrow before distal tibial bone marrow (Fig. 5A). We hypothesized that skull marrow changes may be observable as early as 6 hours following intracisternal injection of 1 × 10^5^ *S. pneumoniae*. Our prior work, which supports this timeline, had identified a rapid skull marrow response in an ischemic stroke model (*4*). We first performed qPCR to clarify if bacteria enter the skull at this early time point and found that the bacterial gene *psaA* was indeed expressed in the skull but not the tibia (Fig. 5B) 6 hours after intracisternal infection. We then analyzed BrdU incorporation into Lin^−^ Sca1^+^ c-kit^+^ hematopoietic progenitors (LSK) in the skull and tibia by flow cytometry to determine whether *S. pneumoniae* had altered LSK proliferation. Mice injected with bacteria had significantly increased BrdU^+^ LSK in their skulls but not in their tibial marrow (Fig. 5C, D). To confirm our observation that elevated LSK proliferation associates with direct local *S. pneumoniae* skull infiltration, we used confocal IVM to image GFP^+^ *S. pneumoniae* progression into the skull marrow 3 hours after injecting bacteria into the cisterna magna. To explore if bacteria were located in the skull marrow’s CSF-containing compartment, we co-injected fluorescently labeled ovalbumin at the time of bacterial infection (Fig. 5E). We noted large extravascular GFP^+^ signal clusters, within the marrow, that were co-labeled with intracisternally injected ovalbumin (Fig. 5F, 5F’), a result suggesting that cells contained bacteria. Additionally, we observed smaller extravascular GFP^+^ areas that also colocalized with the CSF tracer ovalbumin, and we interpreted these as extracellular bacteria (Fig. 5F’’). In sum (Fig. 5G), these data point to a process by which intracisternally injected bacteria co-opt a perivascular CSF passage into the skull marrow, inciting a skull-specific increase in LSK proliferation that precedes changes in distal tibial bone marrow.

**Fig. 5.**
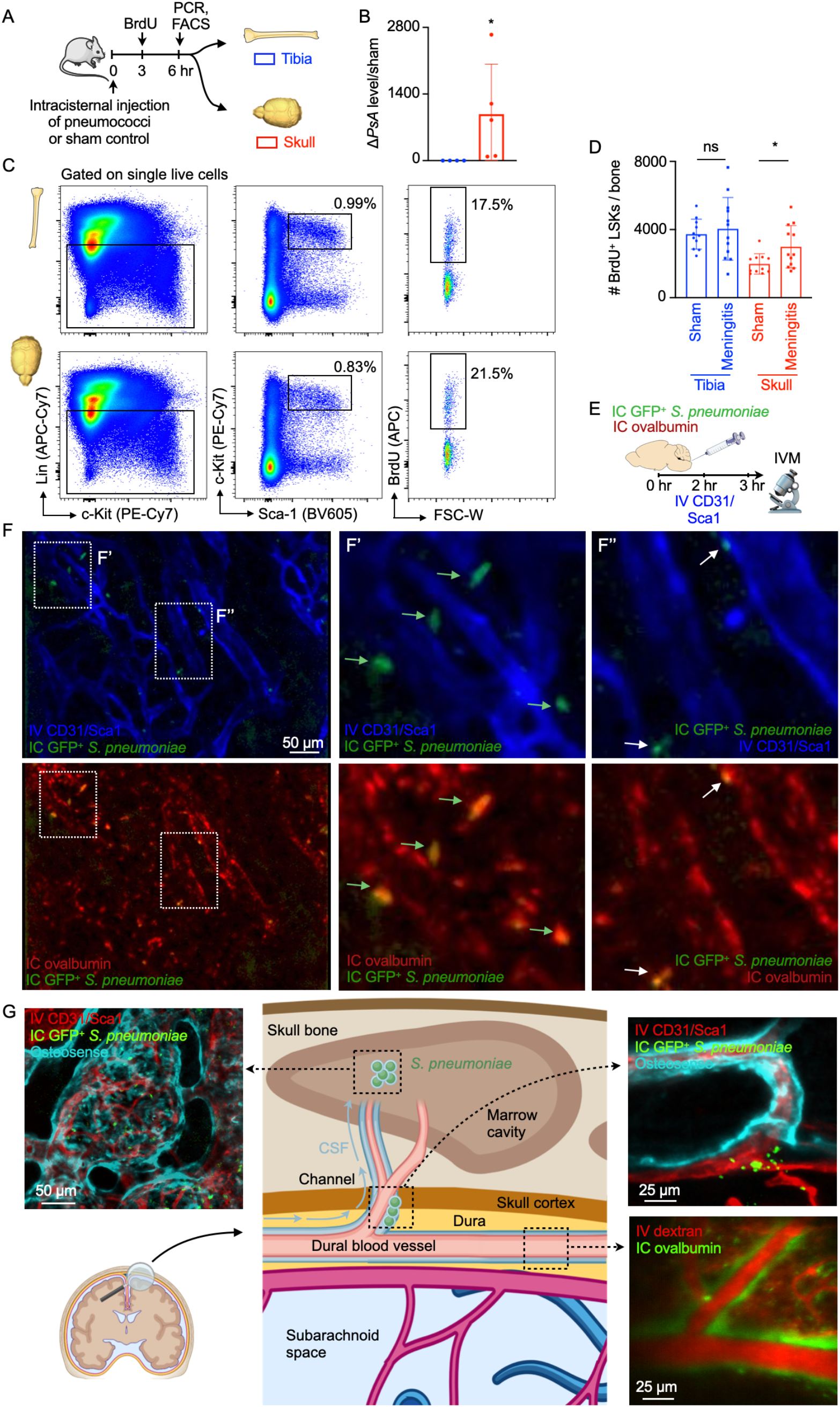
Bacterial meningitis induces LSK proliferation in the skull. (**A**) Outline for experiments (B-D). (**B**) qPCR detection of *S. pneumoniae psaA* gene expression in tibia versus skull normalized to sham controls (mean ± SD; n=4 mice; Mann-Whitney test; \**P*<.05). (**C**) Flow cytometry gating of tibia and skull bone marrow. (**D**) Quantitation of BrdU^+^ lineage^−^ Sca-1^+^ c-kit^+^ hematopoietic progenitors (mean ± SD; n=11-12 mice per group; unpaired t-test; ns=not significant, \**P*<.05). (**E**) Experimental outline. (**F**) IVM of skull BM in mice injected with GFP^+^ *Streptococcus pneumoniae* JWV500 to interrogate bacterial propagation 4-6 hours after intracisternal injection. Bone marrow vasculature was visualized after intravenous injection of fluorescently labeled antibodies for CD31 and Sca1 and CSF outflow by intracisternal injection of ovalbumin. F’ inset depicts large GFP^+^ bacterial areas (arrows). F’’ depicts smaller GFP^+^ areas, presumably bacterial colonies (n=2 mice). (**G**) Summary cartoon indicates CSF outflow via dural perivascular spaces through skull channels into the cranial marrow; this route is usurped by bacteria during meningitis.

## Discussion

Every day, the human ventricular choroid plexus produces most of the 500 ml CSF that provides a protective environment for the brain and receives waste from brain interstitial fluid through exchange with the glymphatic system. CSF outflow is thought to occur via several routes (*2*): i) through arachnoidal villi described more than one hundred years ago (*16*), ii) along spinal and cranial nerves and iii) through dural lymphatics to cervical lymph nodes (*17, 18*). Our work establishes a previously unrecognized outward signaling pathway facilitated by CSF exit along dural vessels that transit skull channels into the marrow.

Given that skull channel CSF outflow — like dural lymphatic vessels — connects to a site of leukocyte abundance, it likely also serves immune surveillance. We speculate that while dural lymphatics alert adaptive immune cells residing in lymph nodes, skull channel signaling may primarily involve innate immune cells produced locally in the calvarial marrow. This reasoning is supported by reports of innate immune cell migration through skull channels in steady state, nerve crush injury, experimental autoimmune encephalomyelitis (*6*) (a mouse model of multiple sclerosis) and chemically induced meningitis (*4*). In contrast to meningeal lymphatics, skull channels transport CSF in a perivascular fashion and facilitate bidirectional exchange. Moreover, skull channels may target spatially distinct areas within meninges, brain and spinal cord because there are many channels that distribute widely within the skull and vertebrae. When considered together, CSF outflow to skull marrow and leukocyte migration towards the meninges appear to be components of a local immune cell supply chain that bypasses systemic circulation, another important distinction from meningeal lymphatics.

During meningitis, the skull marrow is a prominent provider of neutrophils, which reach the meninges through skull channels (*4*). Neutrophils are early responders that combat bacteria in the meninges but also inflict irreversible nerve damage, a common long-term problem in meningitis survivors (*19*). Better understanding skull hematopoiesis as well as neutrophil supply, function and subsets may provide opportunities to support bacterial clearance while alleviating inflammatory CNS damage. The presence of live extracellular bacteria in the skull marrow suggests this hematopoietic tissue may indeed be infected. Intracellular bacteria, on the other hand, may have been removed from the CNS by phagocytes, cells capable of antigen presentation. Bacterial meningitis is most prevalent in newborns and the elderly (*19*), age ranges associated with decreased skull marrow function. The skull is not yet fully developed in neonates, and skull marrow undergoes fatty replacement in advanced age (*20, 21*). We therefore speculate that the bacterial meningitis incidence peaks in those age groups could be related to not only the immune system’s changing competency but also lower hematopoietic capability of the skull marrow. This notion is supported by our observation that hematopoiesis expansion in the cranium outpaced the remote tibial response. Important open questions include whether the phenotype of skull-derived neutrophils differs from cells made in distant marrow regions and how exactly the skull marrow is activated. Prior data on infection-induced emergency hematopoiesis indicate that Toll-like receptors likely sense bacteria (*22*), either directly by hematopoietic progenitors or indirectly via hematopoietic niche cells (Immgen database and deposited data from ref (*23*)).

Our work introduces CSF sampling by the skull marrow, in which immune sentinels are highly abundant. Perhaps related to this discovery, recent human imaging studies showed robust inflammatory signal in skull bone marrow overlying repeatedly abnormal brain cortex in patients with migraine (*24*). While we demonstrate the relevance of CSF outflow to the skull in the setting of bacterial meningitis in mice, such surveillance likely also shapes the immune response in chronic inflammatory CNS disorders such as Alzheimer’s disease and multiple sclerosis. It remains to be determined which cells, cytokines, danger signals, antigens and small molecules traffic through skull channels and how such signaling regulates the production, phenotypes and migration of skull-derived leukocytes towards the CNS. Since dura — marrow connections similar to skull channels exist in vertebrae (*25*), the question arises if vertebral marrow reacts in synchrony with the skull or more like the remote tibial marrow. Generally, the skull marrow warrants closer scrutiny due to its proximity to and crosstalk with the meninges and the CNS.

Constant sampling of CSF outflow suggests the skull marrow state may reflect brain health and that the skull marrow has a prominent role in regulating CNS inflammation.

## Acknowledgements

We acknowledge Jan-Willem Veening for providing fluorescent bacteria and Kaley Joyes for editing the manuscript.

## Author contributions

F.E.P. conceived the study; designed, performed and analyzed imaging and wet lab experiments; induced meningitis; interpreted data and made the figures. J.C.C.-H. designed, performed and analyzed imaging experiments. C.Y. and Z.K. optimized the meningitis model for imaging assays; performed and analyzed experiments; interpreted data and discussed strategy. G.W., D.C., M.Y, J.G., M.J.S. and D. Ro. performed experiments and collected data. C.V. participated in optical clearing and imaging experiments. D.Ri. and J.W.W. provided image analysis. D.B., C.V., D.E.K., F.K.S. and R.W. discussed data and experimental design. F.E.P. and M.N. wrote the manuscript with input from all authors. C.L., M.A.M. and M.N. conceived and directed the study.

## Materials and Methods

### Mice

Mice were housed under certified pathogen-free conditions at Massachusetts General Hospital’s Center for Comparative Medicine. All experiments were conducted in accordance with the Institutional Animal Care and Use Committee’s approval (Protocols: 2005N000306, 2019N000203, 2009N000085 and 2007N000148). Male and female C57BL/6 (WT; JAX 000664) age 10-16 weeks were used for all experiments. Mice were provided rodent chow and water and allowed to acclimate for 1 week before experimentation. All mice were group-housed with *ad libitum* access to food and water. Animals were assigned to experimental groups randomly, and experiments were performed in a blinded fashion.

### Streptococcus pneumoniae

*Streptococcus pneumoniae* strain JWV500 (D39 hlpA-gfp_Cam^r^; serotype 2) was generously provided by Dr. Jan-Willem Veening and prepared as previously described (*26*). *Streptococcus pneumoniae* Xen10 (serotype 3) was purchased from Caliper LifeSciences. GFP^+^ *S. pneumoniae* and *S. pneumoniae* Xen10 were cultured in brain heart infusion broth (BHI) containing 4.5 μg/ml chloramphenicol or 200 μg/ml kanamycin, respectively. *Streptococcus pneumoniae* strains were cryopreserved in BHI with 30% glycerol, thawed the day before the experiment in a 37°C water bath and diluted in fresh BHI with the antibiotic. Bacterial doses between 1:10^3^ and 1:10^6^ CFUs were used, depending on experimental design. Optical densities (600 nm) were used for all bacterial experiments to approximate CFUs, but serial plating dilutions confirmed actual CFUs on BHI agar.

### MicroCT protocol and channel analysis

Samples were imaged using a high-resolution CT scanner (μCT40, Scanco Medical AG, Brüttisellen, Switzerland). Data were acquired using a 6 μm^3^ isotropic voxel size, 70 kVp peak X-ray tube intensity, 114 μA X-ray tube current and 200 ms integration time, and they were subjected to Gaussian filtration. Image renderings were done in Amira (FEI, Hillsboro, OR). Prior to skull channel quantification in Volocity, DICOM files were converted into ND2 file format. Three regions of interest in the midline from the right frontal, parietal and occipital bones were identified and analyzed. Skull channels within regions of interest were visualized on an angled coronal view. Individual channels were given unique identifiers followed by length and width measurements for each channel.

### Cisterna magna injections

Mice were anesthetized by isoflurane inhalation (3-4% induction, 1-2% maintenance), injected with buprenorphine (0.1 mg/kg before surgery and every 12 hrs until sacrifice), followed by hair removal from the back of the neck (Nair). We used a thermometer and feedback-controlled heating blanket (Harvard Apparatus) to maintain body temperature at 37 °C. Mice were fixed on a stereotaxic frame (Harvard Apparatus) with the neck angled downward for optimal cisterna magna exposure, and eye ointment (Dechra) was applied to prevent indirect damage during surgery. An approximately 4 mm vertical skin incision was made at the back of the neck, and the junction between the skull and the 1^st^ vertebrae was exposed by bluntly separating the overlaying muscles (*10*). A 30 μm inner diameter glass micropipette (Fivephoton Biochemicals MGM-1C-30-30) attached to a ultra-precise micro manipulator (Stoelting) loaded with 5 μl of artificial CSF (119 mM NaCl, 26.2 mM NaHCO_3_, 2.5 mM KCl, 1 mM NaH_2_PO_4_, 1.3 mM MgCl_2_, 2.5 mM CaCl_2_) containing 5 × 10^3^ or 1 × 10^5^ *Streptococcus pneumoniae* was inserted through the dura towards the cerebellopontine angle. For sham controls, 5 μl of sterile artificial CSF was injected. These injections were administered at a speed of 1 μl/min with a micro syringe pump (Harvard Apparatus) and a custom-order Hamilton Company syringe (10 μl volume; 3-point style; 20 gauge; 10 mm needle length). In order to prevent backflow-induced variability among individual injections, the needle was retracted incrementally over the course of 10-30 minutes. After the injection, the incision and surrounding area were treated with Terramycin ointment (Zoetis). The incision was sutured with a 5-0 silk suture (Ethilon).

Surgical preparation for 2-photon IVM (described below) was similarly performed with slight modifications to minimize pain and distress over longer imaging periods. Mice were anesthetized with 3% isoflurane, placed on a stereotactic surgery frame (Harvard Apparatus) and then maintained on 1.5% isoflurane in 100% oxygen. Mice were also treated with 0.05 mg per 100 g body weight of glycopyrrolate (Baxter, Inc.), 0.025 mg per 100 g body weight of dexamethasone (07-808-8194, Phoenix Pharm, Inc.) and 0.5 mg per 100 g of ketoprofen (Zoetis, Inc.). Glycopyrrolate and ketoprofen were injected intramuscularly, while dexamethasone was injected subcutaneously. Bupivacaine (0.1 ml, 0.125%; Hospira, Inc.) was subcutaneously administered at the incision site to provide a local nerve block. Animals were provided with 1 ml per 100 g of 5% (w/v) glucose in normal saline subcutaneously every hour during the procedure. We used a thermometer and feedback-controlled heating blanket (40-90-8D DC, FHC) to maintain body temperature at 37 °C. The head and neck were shaved and washed with 70% (v/v) ethanol and iodine solution (AgriLabs). An intracisternal injection was performed as described above. After the injections, the exposed tissue was sealed with cyanoacrylate adhesive (Loctite) and dental cement (Co-Oral-Ite Dental).

### Skull marrow preparation and confocal microscopy

For *in vivo* imaging, the mouse head was shaved and held in a stereotactic skull holder (Harvard Apparatus). Calvarial bone marrow was exposed, as previously described (*11*), by incising a skin flap and then applying glycerol (Sigma-Aldrich) to prevent tissue drying. Skull marrow microscopy was performed with a single photon confocal microscope (IV 100, Olympus, Tokyo, Japan) equipped with IV10-ASW 01.01.00.05 software (Olympus). A field of view at 4x magnification covers a 2290 μm × 2290 μm area while 20x magnification covers a 458 μm × 458 μm area consisting of 512 × 512 pixels. For *ex vivo* skull preparation and imaging after tissue clearing, mice were humanely sacrificed. Then, the head with brain intact was split longitudinally along the sagittal sinus and fixed for 1 hour in 4% paraformaldehyde, after which tissue was washed and subjected to clearing in RapiClear 1.49 (SunJinLab) for 1-2 hours, depending on sample size. Specimen were then mounted on a custom-built tissue holder with a hanging, mounted coverslip (Harvard Apparatus). As indicated in specific figure panels with regard to the intravenous injection timing (retro-orbital, 100 μl total volume in PBS), mice received the following reagents for IVM and *ex vivo* imaging: 30 μl CD31-AF647 (Biolegend, MEC13.3), 30 μl CD31-PE (Biolegend, clone MEC13.3), 30 μl Sca1-AF647 (Biolegend, clone D7), 30 μl Sca1-PE (Biolegend, clone D7) and 100 μl Osteosense 750EX (Perkin Elmer Cat#NEV10053EX).

### *2-Photon IVM and* ex vivo *imaging*

Mice were anesthetized and placed on a custom stereotactic frame. During imaging, anesthesia was maintained with ~1.5% isoflurane in 100% oxygen, with small adjustments to maintain the respiratory rate at ~1 Hz. To fluorescently label the microvasculature, Texas red dextran (40 μl, 2.5%, molecular weight (MW) = 70,000 kDA, Thermo Fisher Scientific) in saline was injected retro-orbitally immediately before imaging. Three-dimensional data sets of the calvarial bone marrow, meninges, meningeal vasculature and CSF transport were obtained using a custom-built two-photon excitation microscope. Imaging was done using 830 nm, 120 fs pulses from a Ti:Sapphire laser oscillator (Spectra-Physics InSight X3). The laser beam was scanned by polygon scanners (30 frames s–1) and focused into the sample using a 60x water-immersion objective lens for high-resolution imaging (numerical aperture of 1.1, Olympus). The emitted fluorescence was detected on photomultiplier tubes through the following emission filters: 400/60 nm for second harmonic generation (SHG), 525/50 nm for Alexa488/FITC and 605/50 nm for Texas red. Laser scanning and data acquisition were controlled by custom-built software. Stacks of images were spaced at 1 μm axially.

### Cerebrospinal fluid tracing

Mice were injected with fluorescent CSF tracers which were reconstituted in artificial cerebrospinal fluid (aCSF) at a concentration of 0.5%. For 2-photon IVM and 2-photon *ex vivo* experiments, mice were IC injected with 2000kD FITC-Dextran (Thermo Fisher Cat# D7137), 70kD Texas Red-Dextran (Thermo Fisher Cat# D1830) and AF647-Ovalbumin (Invitrogen Cat# O34784). For confocal microscopy, mice were injected with AF488-Ovalbumin (Sigma Cat# O34781) and AF647-Ovalbumin (Sigma Cat# O34784) reconstituted in a 5 μl volume of aCSF. We examined CSF trafficking in the marrow by *ex vivo* imaging as described recently(*4*). Z-stacks of marrow-channnel-dural spaces were acquired *in vivo*, after which mice were euthanized at the indicated time points following intracisternal injection of tracer. A piece of frontal bone containing marrow was then excised, preserving the integrity of the dura and the bone marrow cavities. The excised specimen was inverted and rapidly transferred into an aCSF bath or stage-mounted for whole-mount imaging. Tibia bone was embedded in OCT medium (TissueTek) and snap-frozen. Cortical bone was shaved on a crystotat until the marrow was visible.

### Transmission electron microscopy

Skull samples were collected for TEM from naive mice as described above with the following modifications. Cardiac perfusion with PBS was followed by 20 ml of Karnovsky’s fixative (0.1M sodium cacodylate, 2.5% glutaraldehyde, 2% paraformaldehyde). The skull was excised and trimmed to contain the intact frontal, parietal and occipital bones and fixed for 3 hrs in Karnovsky’s fixative followed by 48 hr fixation at 4° C. Skulls were then trimmed to marrow-containing 2 × 4 mm pieces and decalcified over 2 weeks in 140 mM EDTA (pH 7.4; Boston BioProducts). Decalcification solution was replaced with fresh solution every other day and on the final day samples were washed in 0.1M sodium cacodylate buffer for TEM preparation.

After several washes in PBS, followed by several rinses in cacodylate buffer, specimens were fixed in 1% glutaraldehyde in cacodylate buffer overnight at 4°C. The following day, specimens were rinsed several times with cacodylate buffer, infiltrated 1hr in 1% osmium tetroxide, rinsed several times again in cacodylate buffer and then dehydrated through a graded series of ethanols to 100%. Samples were dehydrated briefly in 100% propylene oxide, then incubated in a 1:1 mix of propylene oxide and Eponate resin (Ted Pella, Redding, CA) overnight at room temperature on a gentle rotator. The following day, specimens were incubated at least 3 hr in 100% Eponate resin, then placed into flat molds with fresh 100% Eponate resin and allowed to polymerize in a 60°C oven (24-48hrs). Semi-thin sections (1 μm) were collected onto slides and stained with 0.1% toluidine blue (in 0.1% sodium borate) to preview and confirm the presence of channels. Thin (70nm) sections were cut using a Leica EM UC7 ultramicrotome, collected onto formvar-coated grids, stained with 2% uranyl acetate and Reynold’s lead citrate and examined in a JEOL JEM 1011 transmission electron microscope at 80 kV. Images were collected using an AMT digital imaging system with proprietary image capture software (Advanced Microscopy Techniques, Danvers, MA).

### Clear unobstructed brain/body imaging cocktail and computational analysis (CUBIC)

This protocol was adapted from Tainaka and colleagues (*15*) and used to detect GFP expressed by *S. pneumoniae* in skull channels. Mice were first intracisternally injected with GFP^+^ *S. pneumoniae* as described above for analysis 48 hours after infection. One hour prior to sacrifice, mice were retro-orbitally injected with CD31/Sca1-AF647 cocktail to label the vasculature. Mice were perfused with PBS followed by 4% PFA prior to skull removal. Skull bones were fixed overnight in 4% PFA with shaking at 4°C and subsequently washed 5x for 10 minutes/wash in PBS on a bench-top shaker (400 RPM). Skull samples were delipidized/ decolored in CUBIC-L solution (10 wt% N-butyldiethanolamine (Tokyo Chemical Industry CU#0414), 10 wt% Triton X-100(Sigma)) for 4 days while rotating at 37°C. CUBIC-L solution was refreshed on day 3, and on the final day samples were washed 5x with PBS. Skulls were decalcified over the course of 5 days in CUBIC-B solution (10 wt% EDTA (Boston BioProducts), 15 wt% imidazole (Tokyo Chemical Industry CU#1352)) at 37°C while rotating. Solution was refreshed on day 3, and on the final day samples were washed with PBS. Skull samples were then immersed in CUBIC-L for 2 days at 37°C while rotating and then washed with PBS prior to immunostaining steps. Primary antibody staining using chicken anti-GFP (Abcam, ab13790, 1:300) was performed in staining buffer comprised of PBS with 1% triton-x (Sigma), 10% normal goat serum (Vector Labs) and 0.2% sodium azide (Sigma) for 4 days while gently shaking (200 RPM) at room temperature. Samples were then washed with CUBIC wash buffer (PBS with 1% triton-x) at room temperature 3x for 1 hour per wash after which samples were incubated in goat anti-chicken AF555 (ThermoFisher, A-21437) secondary antibody. This staining was performed in the same buffer used for the primary antibody incubation step at room temperature for 3 days. Samples were washed in PBS 3x for 1 hour per wash and subjected to refractive index matching in CUBIC-R (45 wt% antipyrine (Tokyo Chemical Industry CU#0640), 30 wt% nicotinamide (Tokyo Chemical Industry CU#0855) for 2 days at room temperature, after which samples were ready for whole-mount confocal imaging.

### Bioluminescence imaging

Studies utilizing *S. pneumoniae* Xen10 followed the intracisternal injection technique described above. Briefly, 5 × 10^3^ *S. pneumoniae* Xen10 or an equivalent volume of aCSF was injected into cohorts of age-matched mice for imaging at 12, 36 and 48 hours following injection to track the development of meningitis. Images were acquired with an AmiX BLI/X-Ray Scanner (AmiX) using medium binning and a 3 min exposure time across all time points. Photon intensity was scaled at a range of 4 × 10^4^ – 2 × 10^7^ photons/cm^2^ to allow for cross-group comparisons. The signal was quantified with AMIView software by defining a region of interest across the head, neck and spine. This region of interest was then uniformly fitted to each individual mouse.

### Bacterial colony forming unit assay

CFU assays from blood and CSF of sham and meningitis mice were performed 48 hours after injection. Mice were fixed onto a stereotactic frame in a manner similar to the orientation used for intracisternal injections. To sample the CSF, the injection site was reopened and the dura mater punctured with a glass micropipette to aspirate 5-10 μl CSF. Mice were then removed from the frame and approximately 100 μl of blood was collected by cardiac puncture using a 23 gauge needle and syringe pre-rinsed with 2 mM EDTA. For skull and tibia CFUs, bones were aseptically harvested from sham and meningitis groups, and the meninges were dissected from the skull in sterile PBS containing 5% BSA/2 mM EDTA. Dissection was performed under a stereotactic microscope using fine forceps (#5, angled) and angled spring scissors (Fine Science Tools). Pilot experiments were performed to titer dilutions of CSF, blood, skull or tibia homogenate necessary to visualize bacterial growth on blood agar plates containing 50 μg/ml kanamycin (TEKnova Cat#T0194). A cell-spreader was used to evenly distribute homogenates across the plate and the plate was stored in a 37°C for 24 hours prior to analysis. Plates were photographed and colonies quantitated for relative comparisons.

### S. pneumoniae *detection by qPCR*

We analyzed relative amounts of the *S. pneumoniae* gene *psaA* from tibias and skulls excised from sham and meningitis cohorts of adult mice 48 hours after intracisternal injection of aCSF or *S. pneumoniae* Xen10. Injections were performed as described above and bones were aseptically excised from mice. All mice were perfused with 20 mL PBS prior to removal of tibia and skull bones. After bones were excised, the meninges were dissected from the skull bone, after which tibia and skull bones were snap-frozen in liquid nitrogen and stored at −80°C overnight prior to subsequent analysis. DNA isolation protocol and primers used to detect *S. pneumoniae psaA* were adapted from an established protocol (*27*). Briefly, 3 primers were custom synthesized (IDT) *psaA* forward (5’-GCCCTAATAAATTGGAGGATCTAATGA-3’), *psaA* reverse (5’-GACCAGAAGTTGTATCTTTTTTTCCG-3’) and *psaA* probe (5’-HEX-CTAGCACATGCTACAAGAATGATTGCAGAAA GAAA-3’-phosphate) for qPCR-based relative expression analysis. Skull and tibia bones were trimmed to 50 mg and homogenized with fine scissors in 180 μl ALT buffer containing 0.04 g/ml lysozyme (Sigma) and 75 U/ml of mutanolysin (Sigma). Digestion was performed for 1 hour at 37°C in a shaking water bath. Subsequent steps for DNA isolation and purification were performed following manufacturer guidelines from the Qiagen DNA Mini-Kit manual (Qiagen). gDNA for each sample was spectrophotometrically measured (ThermoFisher NanoDrop 2000) and equivalent DNA amounts were loaded for subsequent qPCR. As described in Carvalho et al., reactions were allowed to run for 45 thermal cycles for amplification and in duplicate for all samples analyzed. Cycle threshold values above 40 were considered negative.

### Flow cytometry

To assess skull and tibia bone marrow hematopoietic cells, mice were anesthetized, sacrificed and perfused with 20 mL PBS to remove blood cells. Tibia and skull were excised, and then meninges were removed from the skull and mechanically homogenized in homogenization buffer (PBS with 5% BSA and 2 mM EDTA). Homogenate was filtered through a 40 μm strainer, centrifuged for 5’ at 340g and resuspended in FACS buffer (PBS with 0.5% BSA) for surface antibody staining. To analyze hematopoietic stem and progenitor cells, cells were first stained with biotin-conjugated anti-mouse antibodies against CD3 (BioLegend, clone 145-2C11), CD4 (BioLegend, clone GK1.5), CD8a (BioLegend, clone 53-6.7), CD49b (BioLegend, clone DX5), CD90.2 (BioLegend, clone 30-H12), CD19 (BioLegend, clone 6D5), B220 (BioLegend, clone RA3-6B2), NK1.1 (BioLegend, clone PK136), TER119 (BioLegend, clone TER-119), CD11b (BioLegend, clone M1/70), CD11c (BioLegend, clone N418) and Gr1 (BioLegend, clone RB6-8C5 all diluted 1:300), which all served as lineage (Lin) markers, and LIVE/DEAD Fixable Aqua Dead Cell Stain (Life Technologies, 1:300). Staining was done for 30 minutes on ice followed by a wash/spin and resuspension in FACs buffer for secondary staining. LSK analysis was performed by staining cells with ckit-PE-Cy7 (BioLegend, clone 2B8), Sca1-BV605 (BioLegend, clone D7) and Streptavidin-APC-Cy7 (BioLegend, 1:100). LSK were identified as Lin^−^ c-kit^+^ Sca-1^+^. Antibodies listed in the secondary staining panel were used at 1:100 dilution in a 500 μl single-cell suspension volume for 30 minutes on ice. Cells were further stained with the APC BrdU Flow Kit (552598, BD Biosciences) following the manufacturer’s guidelines for analysis. BrdU was administered via intraperitoneal injection 3 hours before sacrifice to analyze LSK proliferation 6 hours after intracisternal bacteria injection. Cells were then washed with FACS buffer, spun down at 340g for 5 minutes and resuspended in 400 μl of FACS buffer for analysis. Events were recorded on an LSRII flow cytometer and accompanying FACS DIVA 6.1 software (BD Biosciences). Data were analyzed with FlowJo 10 software (Becton Dickinson).

### Flow cytometric detection of bacteria

Mice were intracisternally injected with GFP+ *S. pneumoniae* as described above and analyzed 48 hours after injection. Skulls were aseptically excised followed by meningeal dissection under a light microscope, as previously described. Samples were then mechanically homogenized in homogenization buffer (PBS with 5% BSA and 2 mM EDTA), filtered through a 40 μm strainer and spun down for 5 min at 340 g. Supernatant containing bacteria was isolated and further spun at 5000 g for 10 min and resuspended in 400 μl to analyze free-floating bacteria. Cell pellets were stained with CD45-AF700 (Biolegend, clone 30F11, 1:300) in FACS buffer on ice for 30 minutes. Samples were washed with 2 ml FACS buffer, spun down at 340 g for 5 min and resuspended in FACS buffer for analysis. Events were recorded on an LSRII flow cytometer and accompanying FACS DIVA 6.1 software (BD Biosciences). Data were analyzed with FlowJo 10 software (Becton Dickinson). Cells were gated for single cells by FSC and SSC parameters, as previously described, and leukocytes were defined as CD45+ cells. GFP+CD45+ cells were discerned for GFP-CD45+ cells, and mean fluorescence intensity was quantitated across groups.

### RNA extraction and qPCR

RNA was isolated from meninges using the RNeasy Micro kit (Qiagen). High-Capacity RNA to cDNA kit (Applied Biosystems) was used for first-strand cDNA synthesis from meningeal RNA. TaqMan gene expression kits were utilized to measure target genes of interest: *Il-1β* (Mm00434228_m1, ThermoFisher), *Il-6* (Mm00446190_m1, ThermoFisher) and *TNF-α* (Mm00443258_m1, ThermoFisher). Target gene primers were all FAM-MGB and all target gene relative expression analyses were normalized to a housekeeping gene, *Gapdh* (VIC-MGB, Mm99999915_g1, ThermoFisher). Samples were run on a 7500 Real-Time PCR machine (Applied Biosystems).

### Image analysis

Images were processed and analyzed using FIJI version 2.1.0, MATLAB R2015b (Mathworks) or Volocity 3D imaging software version 6.3 (PerkinElmer). To complete *ex vivo* 2-PM channel analysis for CSF tracer presence, z-stacks were untilted using a MATLAB code developed in-house, such that the skull surface spanned the least number of z depths, i.e. it lay approximately flat. Afterwards, enhanced contrast was performed on each channel (R,G,B) to correct for intensity attenuation due to light absorption in the tissue. The MATLAB version used was MATLAB 2020b. For confocal microscopy images, z-stacks of .75, 1.0 or 2.0 μm/slice were taken depending on the imaging objective (4x, 10x or 20x). These conditions were consistent across all single-photon confocal microscopy images in both *ex vivo* and intravital microscopy experiments. Images were 3D max-intensity or sum-intensity projected using FIJI and background subtracted followed by an automated de-speckling protocol to remove speckle-based noise. Manual thresholding and contrast adjustment were applied uniformly across all samples when necessary. To ensure processing step uniformity, FIJI macros were recorded for the first image processed and then applied to all images from the same cohort. Where indicated, certain images were further analyzed using Volocity software for preparing 3D surface reconstructions. 3D rendering was automatically generated and image contrast, density and brightness manually and uniformly set across all images analyzed. Movies were manually generated in Volocity using snapshots of post-processed datasets and then incorporated into a single movie in AVI format.

### Statistics

All statistical analyses were performed using GraphPad Prism (GraphPad Software v9.2). All quantitative results are reported as the mean ± standard deviation. For single variable comparisons of parametric datasets for two groups, an unpaired t-test was performed. For non-parametric datasets of unpaired data, a Mann-Whitney test was performed. For parametric datasets for multiple groups, a one-way ANOVA was performed with a post-hoc correction for multiple comparisons as indicated in individual legends. For non-parametric datasets for multiple groups, a Kruskal-Wallis test was performed followed by post-hoc correction for multiple comparisons where indicated.

---

**Fig. S1.**
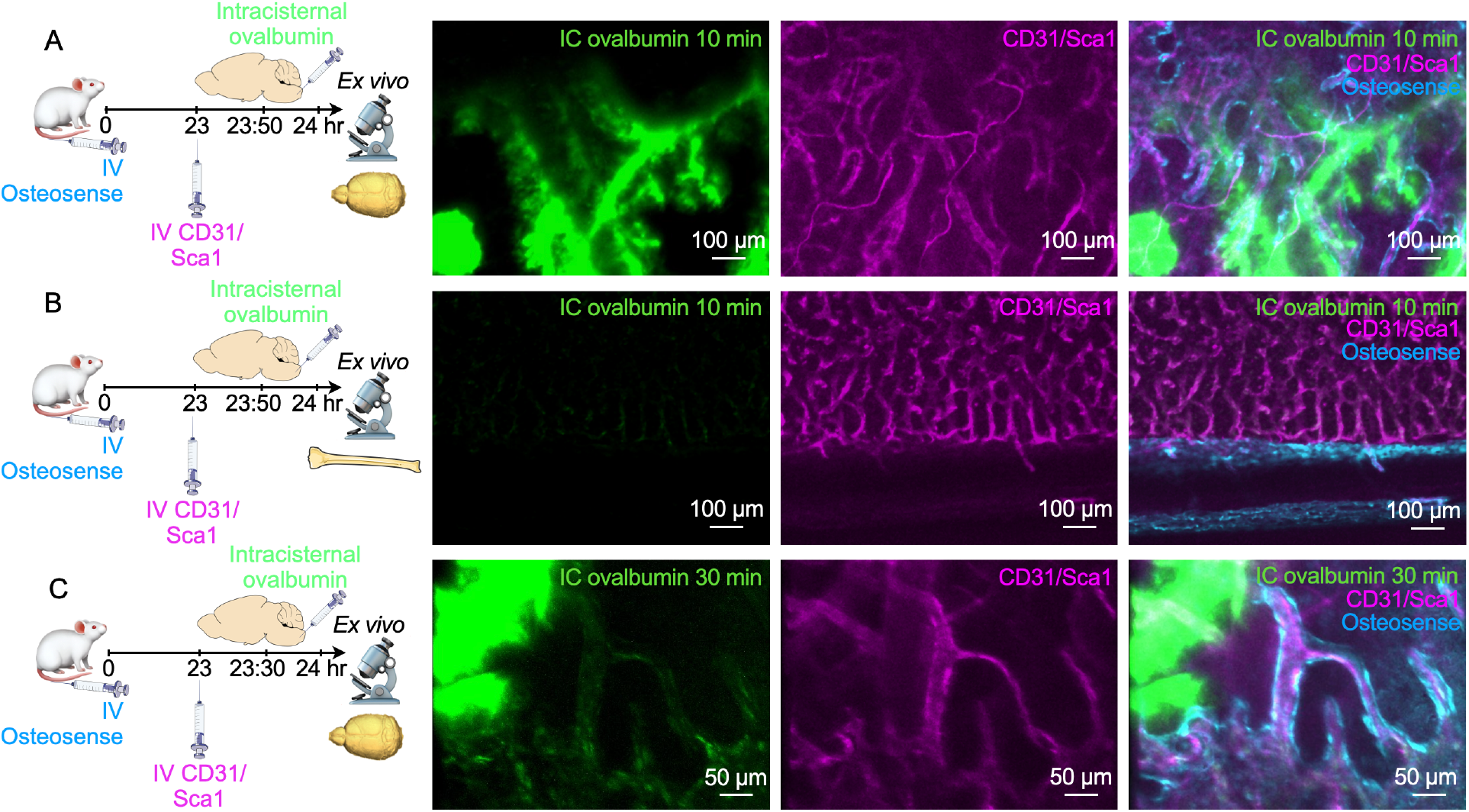
Dynamics of CSF outflow into bone marrow. (**A**) *Ex vivo* imaging of whole-mount skull 10 min after intracisternal (IC) injection of ovalbumin. Intravenous(IV) injection of CD31/ Sca1 labeled vasculature and IV osteosense the bone. (**B**) *Ex vivo* imaging of tibia 10 minutes after intracisternal injection of ovalbumin. (**C**) Imaging 30 minutes after intracisternal injection of ovalbumin.

**Fig. S2.**
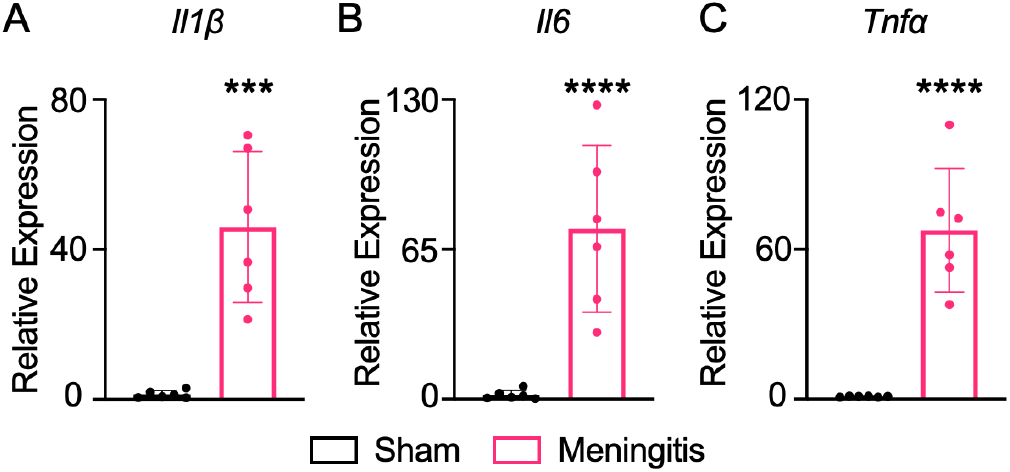
Inflammation in the meninges. qPCR analysis of meninges isolated from either sham controls that were intracisternally injected with artificial CSF or mice 48 hours after intracisternal infection for relative expression analysis of (**A**) *Il1β*, (**B**) *Il6* and (**C**) *Tnfα* (mean ± SD; n=6 mice per group; unpaired t-test; \*\*\*\**P*<.001).

**Movie S1**.

Intravital microscopy of perivascular CSF tracer flow along a dural vessel. Intravenously injected dextran is shown in red, intracisternally injected ovalbumin in green.

**Movie S2**.

*Ex vivo* acquired z-stack of the internal skull cortex showing the CSF tracer (intracisternally injected dextran, green) traveling along channel vessels (intravenously injected dextran, red) into the marrow.

**Movie S3**.

Three-dimensional rendering of an optically cleared skull-brain specimen highlights channels in a sham control mouse. Vasculature was labeled with intravenously injected CD31/Sca-1 antibodies and bone with osteosense.

**Movie S4**.

Three-dimensional rendering of an optically cleared skull-brain specimen from a mouse with meningitis 48 hours after intracisternal injection of GFP^+^ *S. pneumoniae* shows bacterial propagation (green) in the subarachnoid space adjacent to skull channels.

**Movie S5**.

Three-dimensional rendering of a maximal intensity projection acquired by intravital microscopy of skull marrow in a sham control mouse.

**Movie S6**.

Three-dimensional rendering of a maximal intensity projection acquired by intravital microscopy of skull marrow containing GFP^+^ *S. pneumoniae* (green) in a mouse with meningitis.

**Movie S7**.

Three-dimensional rendering of skull channels in a sham control mouse. Vasculature was labeled with intravenous injection of CD31/Sca-1 antibodies and bone with osteosense.

**Movie S8**.

Three-dimensional rendering of a skull channel in a mouse with meningitis shows GFP^+^ *S. pneumoniae* (green) inside the channel. Vasculature was labeled with intravenous injection of CD31/Sca-1 antibodies and bone with osteosense.

